# Selective reactivation of value- and place-dependent information during sharp-wave ripples in the intermediate and dorsal hippocampus

**DOI:** 10.1101/2023.07.12.548282

**Authors:** Seung-Woo Jin, Inah Lee

**Author notes:** Department of Psychiatry and Behavioral Sciences, University of Washington, 1959 N.E. Pacific St, Seattle WA, 98195, USA.

## Abstract

Reactivation of place cells during sharp-wave ripples in the hippocampus is important for memory consolidation. However, whether hippocampal reactivation is affected by the values of events experienced by the animal is largely unknown. Here, we investigated whether place cells in the dorsal (dHP) and intermediate (iHP) hippocampus of rats are differentially reactivated depending on the value associated with a place during the learning of places associated with higher-value rewards in a T-maze. Place cells in the iHP representing the high-value location were reactivated significantly more frequently than those representing the low-value location, characteristics not observed in the dHP. In contrast, the activities of place cells in the dHP coding the routes leading to high-value locations were replayed more than those in the iHP. Our findings suggest that value-based differential reactivation patterns along the septotemporal axis of the hippocampus may play essential roles in optimizing goal-directed spatial learning for maximal reward.

**Teaser:** Information carried by sharp-wave ripples differ qualitatively between the dorsal and intermediate hippocampal regions.

## Introduction

The hippocampus plays a critical role in episodic memory (Eichenbaum 2000, Eichenbaum and Cohen 2001, Squire et al. 2004), which often consists of events associated with different values occurring in space. In the hippocampus, highly synchronized activities of place cells are entrained during sharp-wave ripples (SWRs), and the firing sequences during recent wakeful experiences are re-expressed during SWRs (Wilson and McNaughton 1994). These findings have given rise to the hypothesis that reactivation of place cells contributes to the consolidation process by transferring the newly obtained hippocampal information to neocortical networks (Buzsaki 1989).

It is important to note that prior studies focused mostly on investigating the sequential firing patterns of place cells in the dorsal hippocampus (dHP). For example, it has been demonstrated that an animal’s recent navigational traces or upcoming paths leading to a goal location are reactivated (“replayed”) in the dHP during SWRs (Foster and Wilson 2006, Pfeiffer and Foster 2013). What is largely unknown is whether such reactivations are modulated by different values associated with places. Also, if such value-dependent differential reactivation does occur in the hippocampus, does it occur differentially across different subregions along the longitudinal axis of the hippocampus? A previous study reported that SWRs and replay events occur more frequently in the dHP when a larger reward is provided (Ambrose et al. 2016). Another study showed that the routes associated with larger rewards are replayed more in the dHP than those with smaller rewards (Michon et al. 2019). However, whether such value-based modulation of replay also occurs in other hippocampal regions is still largely unknown.

In our previous study, we found that the place fields of place cells in the iHP were immediately remapped in response to a change in value at a reward location in the T-maze, coalescing around the location associated with the higher-value reward rather than the lower-value reward (Jin and Lee 2021). Other researchers have reported that a subset of place cells in the iHP are dedicated to coding reward information through remapping of their fields to a new place where a reward is available (Gauthier and Tank 2018, Jarzebowski et al. 2022). Therefore, the place cells activated in the dHP and iHP during a navigation task in which places are differentiated by different reward values may also be reactivated differentially during SWRs. We hypothesize that, in the iHP, the neural activities of place cells representing higher-value locations are reactivated more than those representing lower-reward locations, and that such neural firing characteristics may not occur in the dHP, as suggested by our prior study (Jin and Lee 2021).

To test the above hypothesis, we simultaneously recorded neural activities from single units in the dHP and iHP while rats performed a place-preference task on a T-maze (Jin and Lee 2021). We found that place cells in the iHP representing higher-value locations were reactivated more frequently than those coding lower-value locations. However, this value-dependent reactivation pattern was not observed in the dHP. Instead, precise spatial trajectories that sequentially lead to the goal locations were replayed in the dHP, whereas no sequential replay was observed in the iHP. This double dissociation between the iHP and dHP suggests that functionally heterogeneous information content is reactivated during SWRs in different hippocampal regions when goal-directed navigation requires differentiating places based on their values.

## Results

### Behavioral performance in the place-preference task

Rats were trained in a place-preference task on a T-maze (Figure 1A; Video S1). Briefly, in Block 1, rats were required to find the high-value reward arm (HiV-arm) against the low-value reward arm (LoV-arm) by trial and error (Figure 1A) until they reached the learning criterion. To encourage hippocampal-based learning and discourage adopting a response strategy (Packard and McGaugh 1996), in Block 2, rats were started from the opposite side of the original starting point used in Block 1. The spatial relationships between the arms and their reward values were then reversed in Block 3, requiring rats to update the place and its value information (Figure 1A). Spiking activities were recorded from single units in the hippocampus while rats performed the task. Single-unit recordings were also obtained when rats were asleep or immobile before and after the place-preference task (approximately 30 minutes per sleep session).

**Figure 1.**
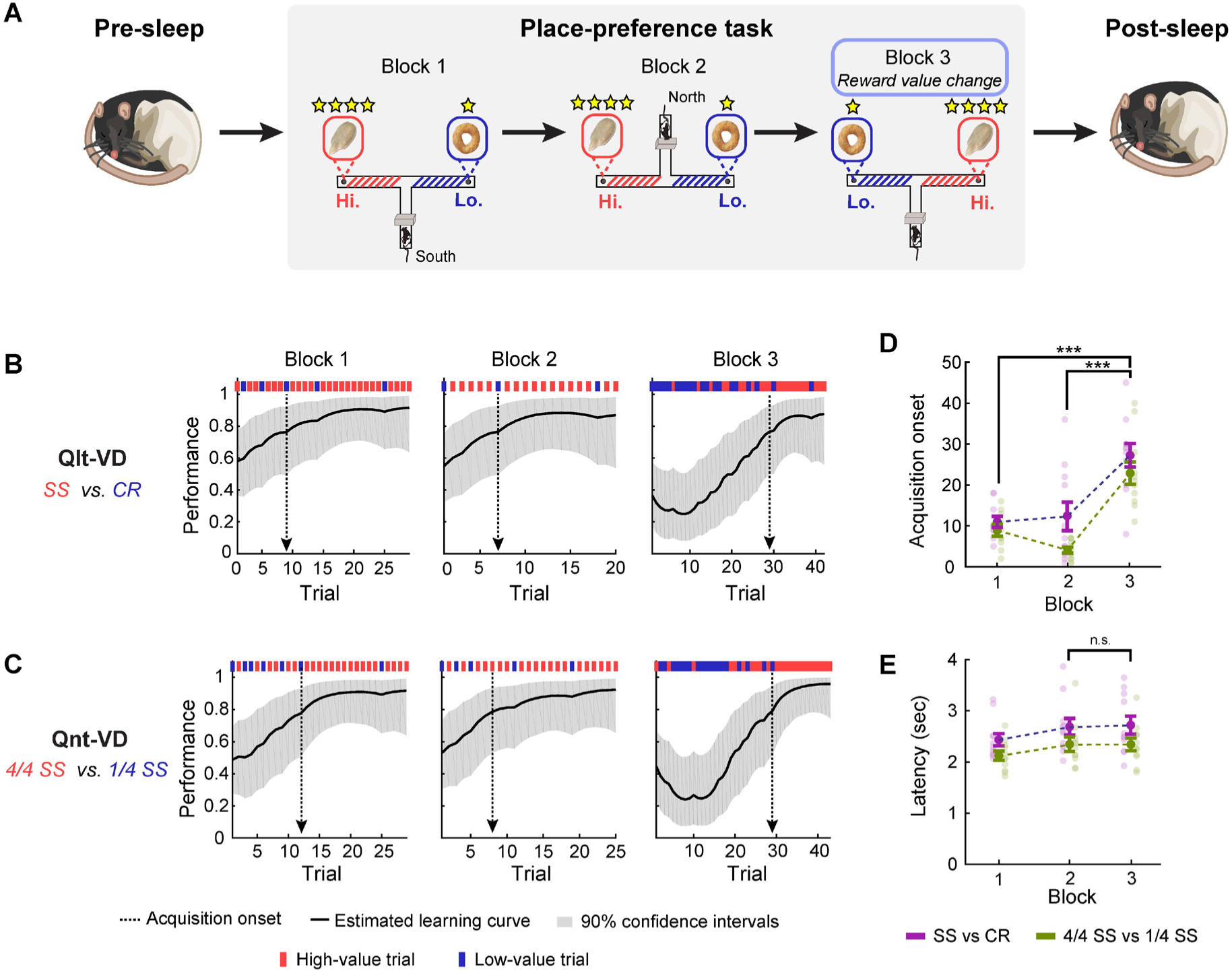
Behavioral performance during the place-preference task. **(A)** Illustration of the place-preference task protocol. Rats started from the end of the south arm or north arm and were isolated at the ends of arms by an acrylic blocker between inter-trial intervals. The arms associated with high-value rewards and low-value rewards were reversed in Block 3 (blue shaded box). **(B** and **C)** Behavioral performance graph of sunflower seed (SS) vs. Cheerios (CR) session (rat 473 – session 2) and 4/4 SS vs. 1/4 SS session (rat 473 – session 3), based on a state-space model (Smith et al. 2004). High-value and low-value trials are marked by red and blue lines, respectively, above the learning curve. The solid black line indicates the estimated learning curve, and the black dashed line indicates the acquisition onset trial, calculated by the state-space model. The 90% confidence interval of the learning curve is shown by gray shading. **(D)** Boxplot for trials that reached acquisition onset for each block. **(E)** Boxplot presentation of mean latency per block. ***p < 0.001.

Reward values were manipulated both qualitatively and quantitatively in our study. Specifically, in sessions 1 and 2 (i.e., *qualitative value-differentiation session*, or *Qlt-VD*), a quarter of a sunflower seed and a quarter of a Cheerios were used as high- and low-value rewards, respectively (Figure 1B). In sessions 3 and 4 (i.e., *quantitative value-differentiation session*, or *Qnt-VD*), a whole sunflower seed and a quarter of a sunflower seed were used as high- and low-value rewards, respectively (Figure 1C). To estimate the trial in which the rat learned the HiV-arm (*acquisition onset*) in each block, we applied a state-space model to behavioral performance data (Smith et al. 2004). Compared to Blocks 1 and 2, learning took place slower in Block 3 (p < 0.001 for both Block 1 and 2 vs. 3 in Figure 1D; Wilcoxon signed-rank test with Bonferroni corrections) (Figure 1D). This difference reflected the fact that rats continually visited the LoV-arm because that arm had been baited with high-value rewards in previous blocks. The delayed onset of acquisition in Block 3 compared to Block 2 cannot be ascribed to a decrease in motivation because their latencies were similar between Blocks 2 and 3 (p > 0.1 for Block 2 vs. 3 in Figure 1E; Wilcoxon signed-rank test). Because the shape of the estimated learning curve and the timing (i.e., trial) of acquisition onset were similar between Qlt-VD and Qnt-VD, neural signals recorded from the two types of sessions were pooled in subsequent analyses.

### More frequent occurrence of reward-zone-associated SWRs in the iHP than in the dHP

Single-unit spiking activities and local field potentials (LFPs) were simultaneously recorded from the dHP and iHP, defined as in our previous study (Jin and Lee, 2021), while rats performed the place-preference task. The SWRs (150-250 Hz) used for analysis were recorded from CA1 pyramidal cell layers in the dHP and iHP (Figure S1A and S1B). SWRs were characterized by large amplitudes (>4 standard deviations [SDs]) and coactivation of at least 5% of cells recorded in a given session while rats showed minimal movement (Figure 2A and 2B; Figure S2A and S2B; see Methods for details). Putative inhibitory interneurons were excluded from subsequent analyses (Figure S1C). Most putative complex spike cells were recorded from the CA1 (Figure S1D).

**Figure 2.**
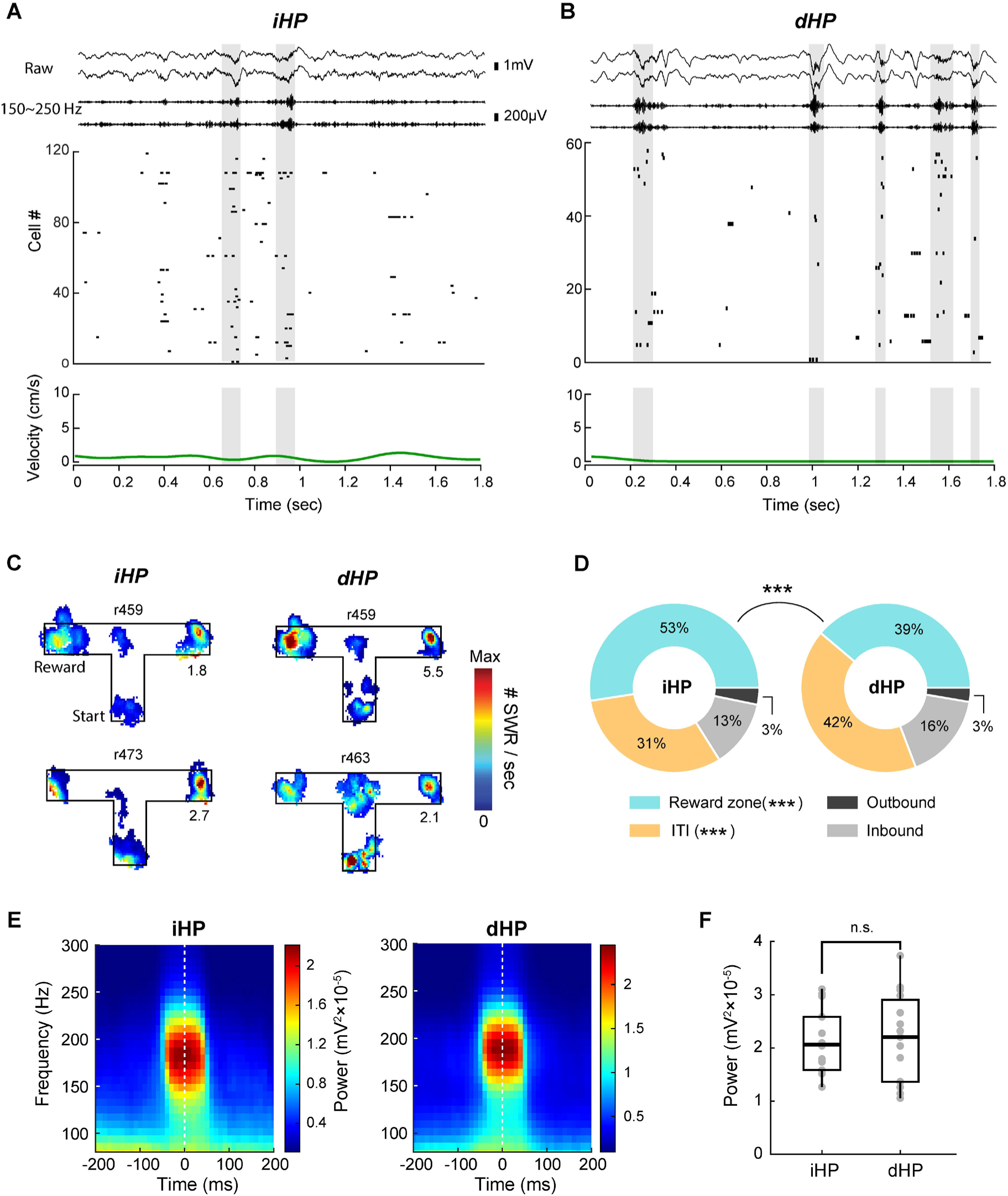
Detection of SWRs during the place-preference task. **(A** and **B)** Graphs of raw LFP, 150–250-Hz filtered signal, ensemble firing, and velocity illustrating the SWR periods that satisfied SWR criteria. SWR periods recorded from the iHP **(A)** and dHP **(B)** are indicated by gray-shaded areas. The scale indicates the amplitude of raw and filtered LFP signals. See also Figure S1. **(C)** Results for two individual rats exemplifying the distribution of SWR occurrence in the iHP and dHP. The black line indicates the contour of the maze; a spatial bin with lower SWR rates was excluded from the color display. SWRs recorded from Blocks 1 and 3 were pooled for calculation of the SWR rate map. See also Figure S2. **(D)** Pie chart showing the frequency of SWRs recorded from the dHP and iHP when rats were in the outbound zone, reward zone, inbound zone, and ITI zone during a place-preference task. **(E)** Spectrogram displaying the frequency and power of SWR, aligning with the time of SWR peak amplitude. **(F)** Comparison of peak amplitude of SWR between the iHP and dHP. ***p < 0.001.

Based on our SWR-detection criteria, a significant majority of SWR events (approximately 80-85%) took place when rats were located in reward zones consuming the reward or during inter-trial interval (ITI) periods while rats waited for the next trial (Figure 2C and 2D; Figure S2A and S2B). Similar proportions of SWRs occurred between the reward zone and ITI period in the dHP (Figure 2D). However, approximately half of SWR events in the iHP occurred while rats were in the reward zone, suggesting that the effect of the reward on the occurrence of SWRs was more pronounced in the iHP compared to the dHP (p < 0.001 for group comparison; p < 0.001 for reward zone and ITI conditions; Chi-square test with Bonferroni corrections) (Figure 2D). Additionally, the properties of SWRs occurring within the reward zones, including their frequency and power, exhibited similar characteristics in both the dHP and iHP (p > 0.1 for power between iHP and dHP; Wilcoxon rank-sum test) (Figure 2E and 2F). Since we are mostly interested in the reward value-related neural reactivation in the dHP and iHP, we restrict our analysis of SWR events to the reward zones, reflecting the high likelihood of a substantial disparity in the proportion of SWR events within these zones between the dHP and iHP.

### Exclusive reactivation of place cells representing high-value locations during correct trials in the iHP but not in the dHP

During the acquisition of reversed reward contingency, some place fields underwent global remapping toward arms associated with either high-value or low-value rewards, indicating a significant influence of reward value on these cell’s spatial representations. On the other hand, certain fields maintained their place representations, irrespective of changes in reward value, suggesting their primary role in encoding spatial information (Figure 3A and 3B). The degree of alteration in spatial firing patterns following the reversal of reward contingency was measured by Pearson’s correlation coefficients between rate maps before and after the reward reversal. When these coefficients fell below 0.75 which was an operational criterion of global remapping, these cells were classified as spatial-value-coding (or value-coding) place cells. Depending on whether their place fields were located within either HiV-arm or LoV-arm, these cells were further categorized as either high-value-coding place cells (HiV-cells) or low-value-coding place cells (LoV-cells). Place cells with stable representations, exhibiting coefficients exceeding 0.75, were designated as location-coding place cells (Location-cells). Place cells with their place fields located in the stem were excluded from the main analysis because it was challenging to determine whether these cells represented high-value or low-value information. The number of HiV-cells and LoV-cells was found to be similar in both the iHP (n = 27 for HiV-cells, n = 26 for LoV-cells; p > 0.1 for one-sample Chi-square test) and dHP (n = 23 for HiV-cells, n = 24 for LoV-cells; p > 0.1), suggesting that value– coding place cells are uniformly distributed in both HiV-and LoV-arms (Figure 3C).

**Figure 3.**
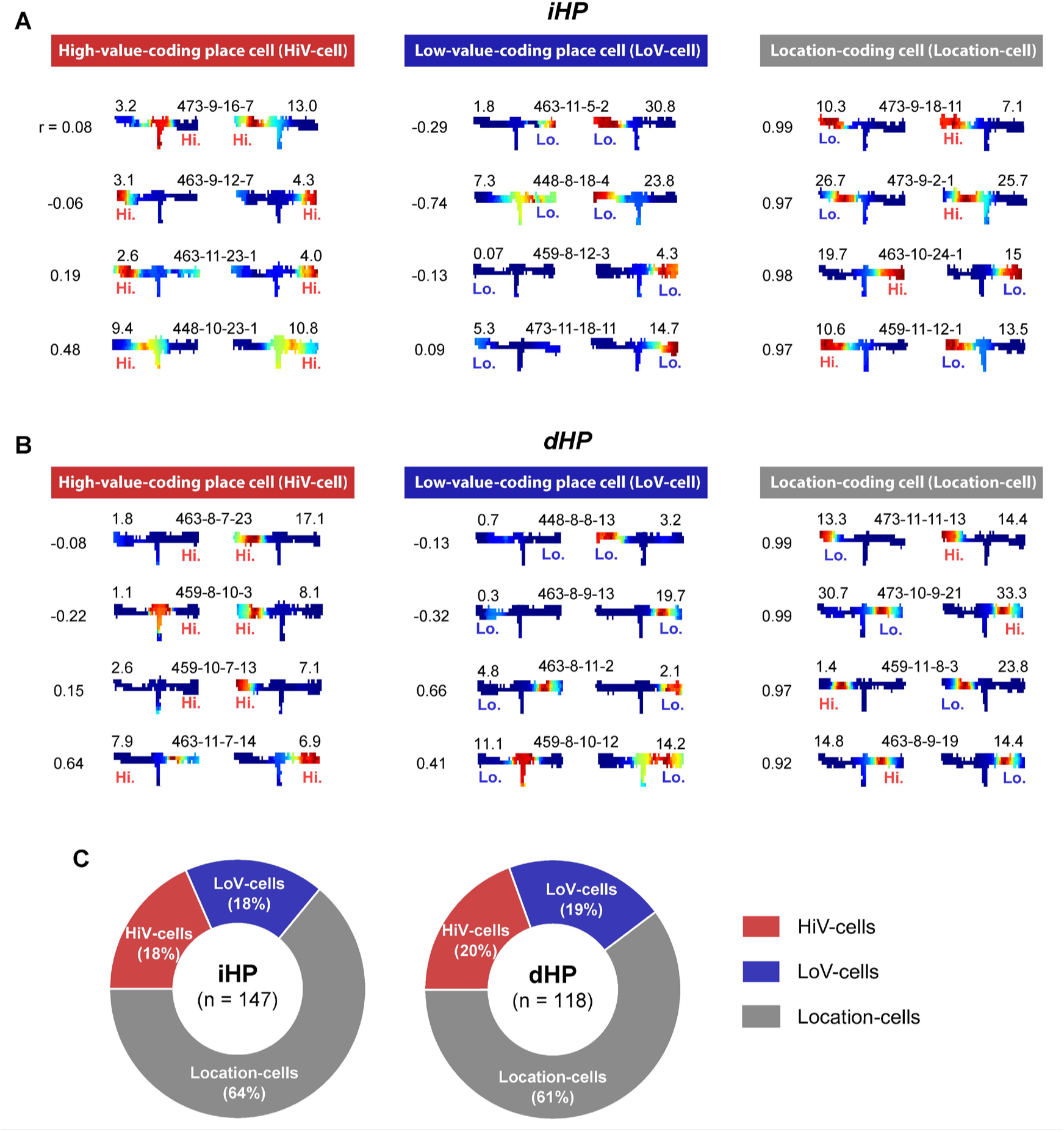
Classification of place cells into three types. **(A** and **B)** The single-cell examples of showing HiV-cell, LoV-cell, and Location-cell. The number on the left side of the rate maps indicates the correlation coefficient between spatial rate maps. Cell identity (rat # – session # – tetrode # – cluster #) and peak firing rates were annotated between rate maps and above each rate map, respectively. **(C)** Pie chart showing the number of HiV-, LoV-, and Location-cells.

Since location- and value-coding place cells were best dissociated in Block 3, in which rats learned the reversal task, SWR-based reactivation patterns were further analyzed only in the reversal block (i.e., Block 3). Our specific aim was to examine whether HiV-cells exhibited a higher reactivation probability compared to LoV-cells or Location-cells. Individual SWR examples revealed that among the three cell types (HiV-, LoV-, and Location-cells), spiking activities of HiV-cells were reactivated exclusively during SWRs in the reversal block in the iHP (Figure 4A). To compare the reactivation probability during the reversal block across these cell types at the population level, we employed a peri-event time histogram aligning all spikes to the moment of the SWR peak. Comparing HiV-cells to LoV-cells and Location-cells, it became evident that HiV-cells had greater reactivation probabilities in the vicinity of the SWR amplitude peak (-40ms ∼ +40ms; periods marked with a thick line above the graph, where p < 0.05 of Kruskal-Wallis test for each time bin) (Figure 4B). Furthermore, single HiV-cells in the iHP had a higher probability of reactivation during SWRs when compared to both LoV-cells (p < 0.01) and Location-cells (p < 0.05; Wilcoxon rank-sum test with Bonferroni corrections) (Figure 4C).

**Figure 4.**
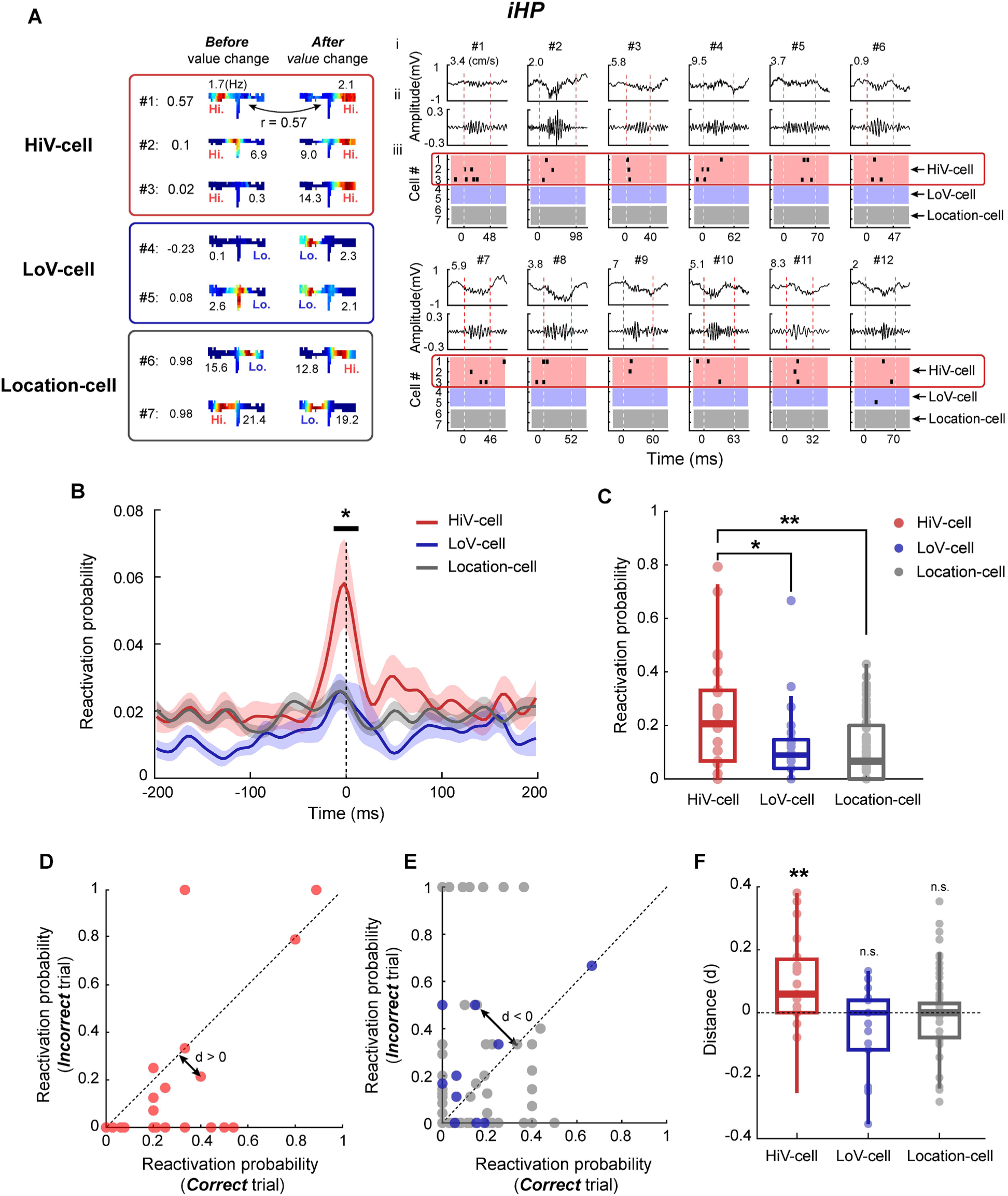
Exclusive reactivation of HiV-cells in the iHP during SWRs. **(A)** Illustrative examples of spatial rate maps for three cell types in the iHP. These cells #1-7 were observed in rat 473 – session 3. Peak firing rates were indicated above the rate maps, and correlation coefficients between rate maps were shown on the left side of the rate maps. Spatial rate maps of seven single cells in the iHP are shown, together with their reactivation patterns during awake SWRs in Block 3. Spatial rate maps for each type are grouped as denoted by colored boxes. Right panel: Examples of SWR and reactivation activities occurring after the reward value was changed. Raw LFP (i), 150–250-Hz filtered signals (ii), and reactivation of Cells #1-7 (iii) per SWR period are shown. Mean velocity during an SWR is shown on the raw LFP graph. The dashed line indicates the boundary of an SWR. **(B)** Peri-event time histogram aligning with the SWR peak time, illustrating the relationship between reactivation probability and SWR events depending on the cell types. **(C)** Comparison of reactivation probabilities in the iHP. Each data point indicates the reactivation probability of a single cell. **(D** and **E)** Scatter plot displaying the distribution of reactivation probability between correct and incorrect trials. The perpendicular distance between the diagonal line and each data point was calculated to assess the degree of bias regarding whether cells exhibited a higher reactivation probability in either correct or incorrect trials. **(F)** Comparison of the distance depending on cell types. Each data point indicates the distance of a single cell. See also Figure S3. *p < 0.05, **p < 0.01, ***p < 0.001.

Furthermore, we analyzed to investigate whether high-value rewards in correct trials and low-value rewards in incorrect trials had distinct effects on exclusive reactivation of HiV-cells. To address this, we separately calculated the reactivation probability of individual cells depending on whether the trials were correct or incorrect. We graphically represented the results using a scatter plot, positioning a data point on the diagonal line if a cell exhibited an equal probability for both correct and incorrect trials. However, if cells displayed a greater reactivation probability in correct trials than in incorrect trials, data points were placed below the diagonal line. Notably, the red data points representing HiV-cells were predominantly distributed below the diagonal line, indicating that these cells tended to exhibit greater reactivation during correct trials than incorrect trials (Figure 4D). In contrast, the distribution of data points for LoV- and Location-cells did not show a consistent bias toward either the upper or lower side of the diagonal line (Figure 4E). To quantitatively assess the extent of deviation from the diagonal line, we computed the perpendicular distance between each data point and the diagonal line. The distances of HiV-cells were significantly higher than 0, while for LoV-cells and Location-cells, the distances did not significantly differ from 0 (p < 0.01 for HiV-cells, p > 0.1 for LoV-cells and Location-cells; One-sample Wilcoxon Signed-rank test with Bonferroni corrections) (Figure 4F).

Conversely, when we applied the same analysis to the dHP as depicted in Figure 4, it became evident that all cell types exhibited similar levels of reactivation probabilities during SWRs (Figure 5A). The peri-event time histogram revealed that HiV-, LoV- and Location-cells displayed comparable reactivation probabilities during the SWR events (Figure 5B). This uniformity in reactivation probabilities in the dHP was further supported at the individual place-cell level, where we observed that probabilities of reactivation during SWRs were similar across all cell types (p > 0.1 for HiV-cells vs. LoV-cells and Location-cells; Wilcoxon rank-sum test with Bonferroni corrections) (Figure 5C). Furthermore, there was no significant difference in reactivation probabilities between correct and incorrect trials (Figure 5D to 5F). This disparity between the dHP and iHP was further confirmed by reactivation rates as an alternative measurement (Figure S3A). Notably, the elevated reactivation probabilities of HiV-cells in the iHP cannot be attributed to the higher excitability of this cell type, as there was no significant correlation between the peak firing rates of individual place cells and their reactivation probabilities (Figure S3B). Therefore, our findings suggest that the exclusive reactivation of place cells representing high-value locations during SWRs is a unique property of the iHP.

**Figure 5.**
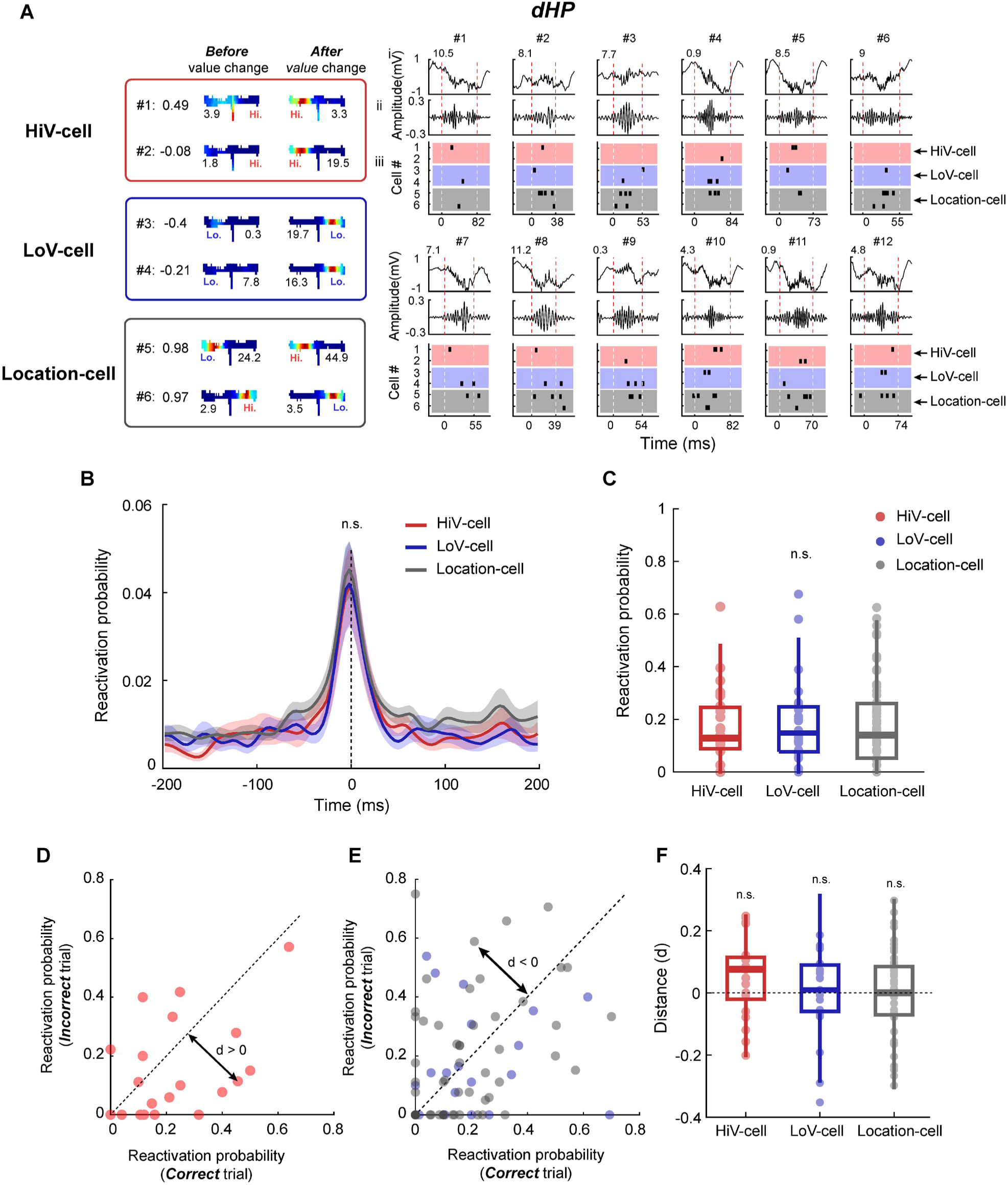
Uniform reactivation of all cell types in the dHP. **(A)** Illustrative examples of spatial rate maps for three cell types in the dHP. These cells #1-6 were observed in rat 463 – session 1. Peak firing rates were indicated above the rate maps, and correlation coefficients between rate maps were shown on the left side of the rate maps. Spatial rate maps of seven single cells in the dHP are shown, together with their reactivation patterns during awake SWRs in Block 3. Spatial rate maps for each type are grouped as denoted by colored boxes. Right panel: Examples of SWR and reactivation activities occurring after the reward value was changed. Raw LFP (i), 150–250-Hz filtered signals (ii), and reactivation of Cells #1-6 (iii) per SWR period are shown. Mean velocity during an SWR is shown on the raw LFP graph. The dashed line indicates the boundary of an SWR. **(B)** Peri-event time histogram aligning with the SWR peak time, illustrating the relationship between reactivation probability and SWR events depending on the cell types. **(C)** Box plot comparing the reactivation probabilities in the dHP. Each data point indicates the reactivation probability of a single cell. **(D** and **E)** A scatter plot displaying the distribution of reactivation probability between correct and incorrect trials. The perpendicular distance between the diagonal line and each data point was calculated to assess the degree of bias regarding whether cells exhibited a higher reactivation probability in either correct or incorrect trials. **(F)** Comparison of the distance depending on cell types. See also Figure S3.

### Learning-dependent potentiation of exclusive reactivation of high-value-coding place cells in the iHP but not in the dHP

Next, we investigated whether the reactivation of HiV-cells during SWRs in the iHP was enhanced as rats acquired the reversed relationships between place and its associated value during Block 3. In the pre-learning phase, HiV-cells exhibited less frequent reactivation during SWRs (left panel in Figure 6A). However, as rats reached the acquisition onset, a notable change in reactivation patterns occurred. HiV-cells in the iHP began to fire exclusively during SWRs, while other cell types remained silent (right panel in Figure 6A). We carried out a thorough investigation into learning-dependent alterations in reactivation by segregating trials into five trial distinct bins based on performance (i.e., < 50%, 50∼65%, 65∼75%, 75∼90%, >90%). In this analysis, we used performance of 75% as an indicator of learning because acquisition onset, as computed by a state-space model, was consistently around 75% (Figure 1B and 1C). The results revealed that once performance exceeded 75%, HiV-cells exhibited an upsurge in reactivation probability, with these cells having a greater reactivation probability than other cell types (75∼90%: p < 0.001 for HiV-cell vs. LoV-cell; > 90%: p < 0.01 for HiV-cell vs. LoV-cell; Kruskal-Wallis test). In contrast, the remaining cell types did not exhibit notable increases in the vicinity of acquisition onset (Figure 6B).

**Figure 6.**
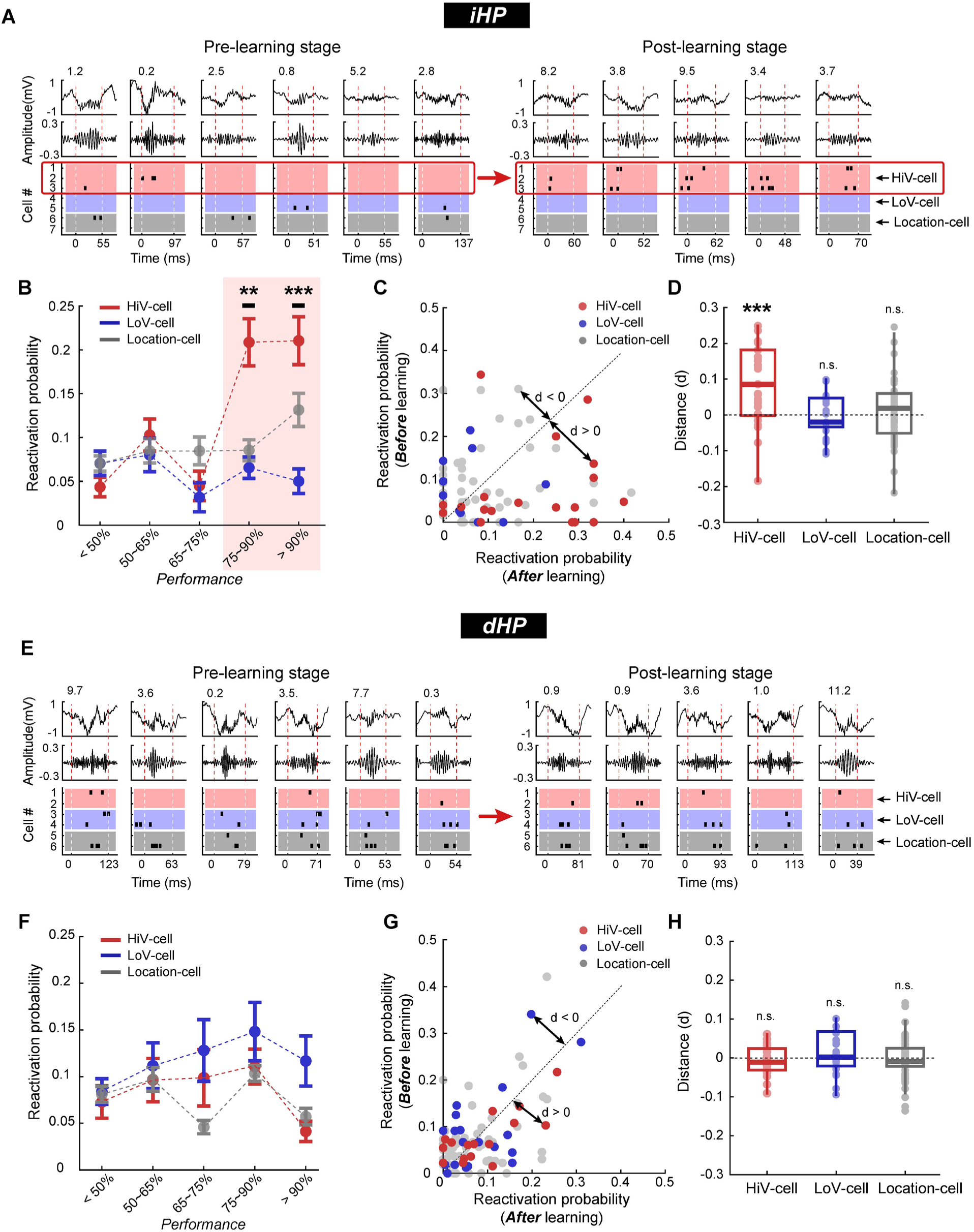
Learning-dependent changes in the reactivation of HiV-cells in the iHP. **(A)** Illustrative examples of awake SWR and reactivation activities in the iHP that occurred during correct and incorrect trials in the pre-learning state, and correct trials in the post-learning state. The cells used in (A) are the same as those in Figure 4. Spatial rate maps for each type are grouped together as denoted by colored boxes. **(B)** The changes in reactivation probability depending on the performance. The correctness was divided into five types. **(C)** Scatter plot displaying the distribution of reactivation probability between pre-learning and post-learning phases. The perpendicular distance between the diagonal line and each data point was calculated to assess the degree of bias regarding whether cells exhibited a higher reactivation probability in either the pre-learning or post-learning phase. **(D)** Comparison of the distance depending on cell types. Each data point indicates the distance of a single cell. (**E** – **H**) The same analyses as in (**A** – **D**) were replicated. See also Figure S4. *p < 0.05, ***p < 0.001.

To evaluate the change in reactivation probability before and after the acquisition of the task, we utilized a scatter plot that displayed reactivation probabilities in the pre-learning and post-learning phases. It was evident that the majority of data points representing HiV-cells (red points) were positioned below the diagonal line, whereas those for LoV-cells (blue points) and Location-cells (gray points) were distributed both above and below the diagonal line (Figure 6C). This pattern indicated that only HiV-cells exhibited an increase in their reactivation probability after learning. For statistical testing, we calculate the distance between each data point and the diagonal line. The distance of HiV-cells was significantly higher than 0, whereas other cell types did not show statistical significance (p < 0.001 for HiV-cells, p > 0.1 for LoV-cells and Location-cells; One-sample Wilcoxon Signed-rank test with Bonferroni corrections) (Figure 6D). These findings strongly support the idea that the exclusive reactivation of HiV-cells in SWRs in the iHP is contingent on the learning process.

In contrast, when the same analysis was applied to the dHP, we did not observe learning-dependent changes in the reactivation of HiV-cells in the dHP, akin to what was observed in the iHP (Figure 6E and 6F). The distance from the diagonal line did not show significant increases or decreases compared to 0 (p > 0.1 for HiV-cells, LoV-cells, and Location-cells; One-sample Wilcoxon Signed-rank test with Bonferroni corrections) (Figure 6G and 6H). Collectively, these results suggest that the reactivation of place cells representing the high-value location upon reversal in the iHP may play a critical role in consolidating the newly learned spatial value.

### Selective reactivation of the place cells representing high-value locations in the iHP during the sleep session after learning is correlated with the next day’s place-value learning

To assess whether the potentiated reactivation of HiV-cells in the iHP persists in the post-sleep session following a behavioral recording session, we examined reactivation patterns during both pre- and post-sleep sessions, using the same place cells as depicted in Figure 4. In the pre-sleep session, HiV-cells in the iHP were rarely reactivated during SWRs (left panel in Figure 7A). However, following the behavioral session of the place-preference task, exclusive reactivation of HiV-cells was observed in the post-sleep session (right panel in Figure 7A). Importantly, those HiV-cells that exhibited a higher reactivation probability during the behavioral session also had a higher reactivation probability measured by Δsleep (i.e., post-sleep – pre-sleep). This observation suggests that potentiated reactivation during behavioral sessions was sustained during the post-sleep session (p < 0.05, r^2^ = 0.20; linear regression) (Figure 7B).

**Figure 7.**
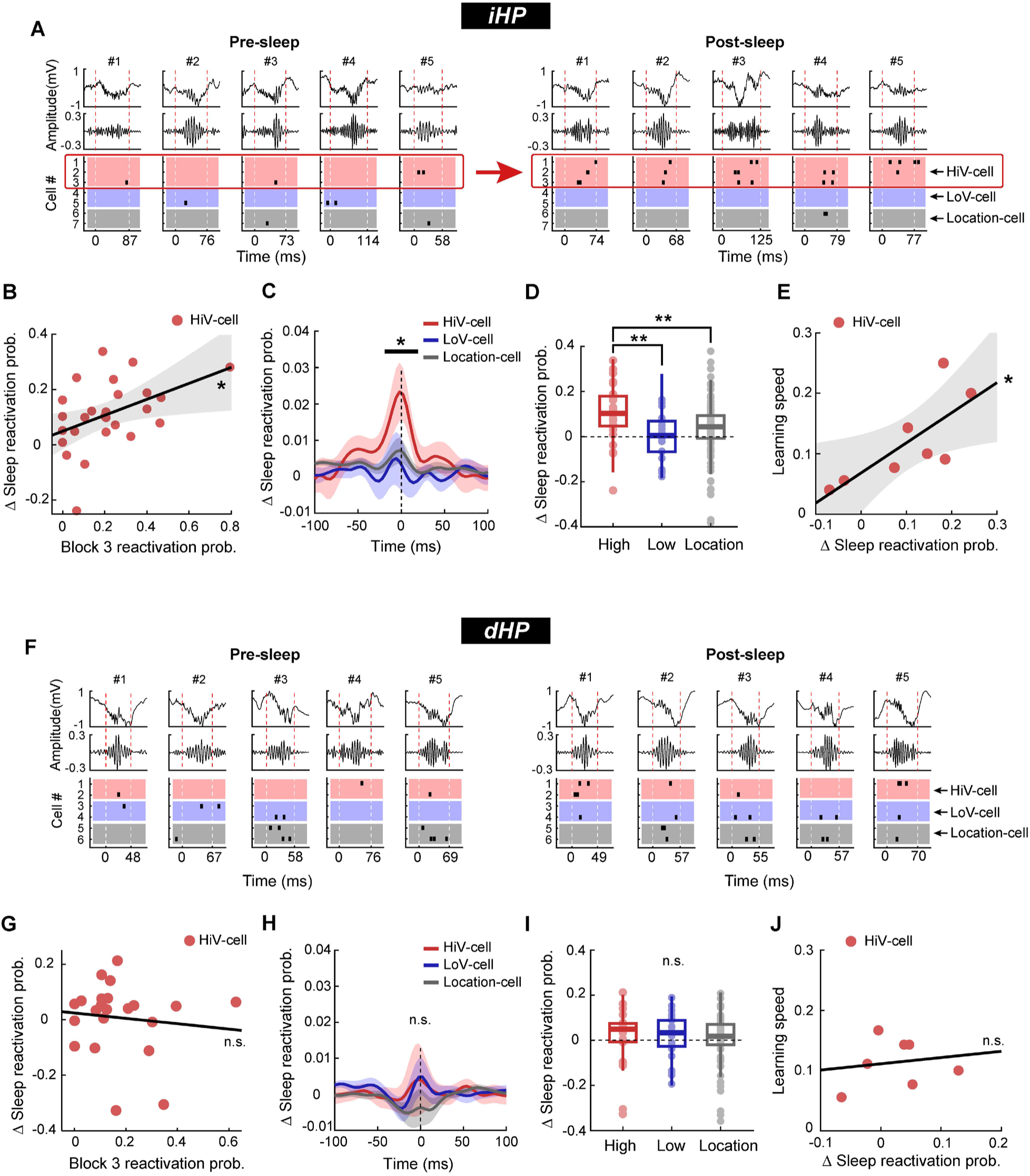
Enhanced reactivation of HiV-cells in the iHP, but not in the dHP, is maintained during sleep after the place-preference task. **(A)** Illustrative Examples of sleep SWR and reactivation activities that occurred during pre-sleep and post-sleep in the iHP. Raw LFP, 150–250-Hz filtered LFP, and reactivation of Cells #1-7 during the SWR period are shown. The cells used in (A) are the same as those in Figure 4. Dashed line indicates the boundary of an SWR. **(B)** Scatter plot illustrating the correlation between reactivation probability during the behavioral session and sleep by computing the degree of correlation between them. Each data point indicates the reactivation probability of a single cell. **(C)** Peri-event time histogram aligning with the SWR peak time, illustrating the relationship between Δsleep reactivation probability and SWR events depending on the cell types. **(D)** Comparison of Δsleep reactivation probabilities in the iHP. Each data point denotes the reactivation probability of a single cell. **(E)** Examination of the relationships between Δsleep reactivation probability and learning speed on the subsequent day, based on the session mean reactivation probability of HiV-cells in the iHP. Each data point represents the average reactivation probabilities of HiV-cells per session. The dotted line indicates the 95% confidential interval of the linear regression. (**F** – **J**) The same analyses as in (**A** – **E**) were replicated in the dHP. See also Figure S6. p < 0.05, **p < 0.01, ***p < 0.001.

To quantitatively compare Δsleep reactivation probability among cell types, we calculated the peri-event time histogram by aligning the time at the peak of SWR amplitude. This analysis revealed that the Δsleep reactivation probability of HiV-cells was significantly greater than those of other cell types, particularly in the vicinity of SWR peak amplitude (-30ms ∼ +30ms; periods marked with a thick line above the graph where p < 0.05 of Kruskal-Wallis test for each time bin) (Figure 7C). Furthermore, when we examined Δsleep reactivation probability for individual cells, we found the probability of HiV-cells was greater than that of other cell types in the iHP (p < 0.01 for HiV-cells vs. LoV-cells and Location-cells; Wilcoxon rank-sum test with Bonferroni corrections) (Figure 7D). Importantly, the increase in Δsleep reactivation probability of HiV-cells in the iHP exhibited a positive correlation with the learning speed (i.e., the inverse of acquisition onset) in Block 1 on the following day (p < 0.05, r^2^ = 0.56; linear regression) (Figure 7E). This result suggests that reactivation of HiV-cells during post-sleep plays a role in consolidating the high-value location, thereby enhancing spatial learning on the subsequent day.

In contrast, in the dHP, there were no detectable changes between pre- and post-sleep sessions (Figure 7F). Specifically, we barely observed a positive correlation between reactivation probabilities during behavioral sessions and sleep (p > 0.1, r^2^ = 0.004; linear regression) (Figure 7G). Additionally, there was no significant difference in Δsleep reactivation probability across cell types at the SWR peak (Figure 7H) (p > 0.1 of Kruskal-Wallis test for all of each time bin), and Δsleep reactivation probability did not significantly differ among cell types (p > 0.1 for Kruskal-Wallis test) (Figure 7I). Furthermore, positive correlations were rarely found between Δsleep reactivation probability and learning speed of the next day in the dHP (p > 0.1, r^2^ = 0.03; linear regression) (Figure 7J). We confirmed these findings by conducting separate analyses for the pre- and post-sleep sessions (Figure S4). Additionally, the positive relationship in the iHP between the amount of Δsleep reactivation probability and learning speed of the next day was not observed in LoV-cell and Location-cell both in the iHP and dHP (Figure S5). Collectively, our findings imply that reactivation of HiV-cells in the iHP during sleep may underlie the consolidation of value-updated location information.

### Sequential spatial reactivation of place cells that lead to the high-value reward location is found only in the dHP

Finally, we examined whether sequential reactivation of place cells (i.e., spatial replay) which has been frequently reported in the dHP in the literature (Pfeiffer and Foster 2013, Diba and Buzsaki 2007, Foster and Wilson 2006), is detected in the iHP. Initially, we selected a route from a start location to a high-value location and visually represented this route with a color-coding scheme according to value (Figure 8A). Population rate maps were constructed by combining place fields of ensemble cells from the dHP and iHP, respectively (Figure 8B). Subsequently, using Bayesian decoding, we calculated the posterior probability for place cells that were recorded simultaneously during a SWR event (Figure 8C and 8D). In the dHP, place cells representing different places from the start location (indicated by blue data points in Figure 8C) to the reward location (marked by red data points in Figure 8C) were sequentially reactivated in either a forward direction (forward replay in #1 – 4 in Figure 8C) or a reverse direction (reverse replay in #5 – 8 in Figure 8C) within individual SWR events.

**Figure 8.**
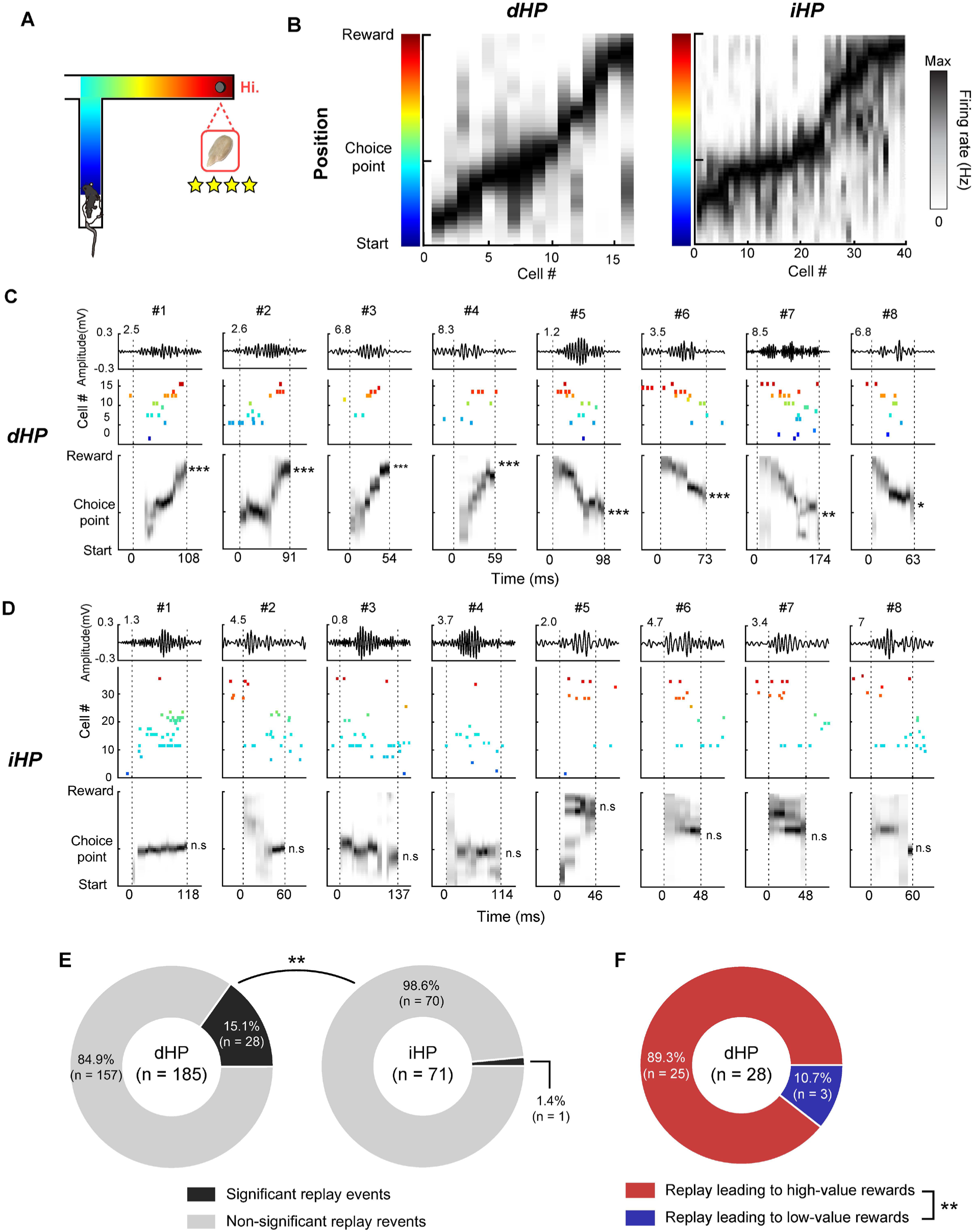
Spatial replay leads towards the goal location in the dHP but not in the iHP. **(A)** T-maze illustration with color-coded positions. The color-coding map was utilized to visualize the spatial replay examples presented in (C) and (D). **(B)** Population rate maps of ensemble place cells constructed from the dHP (left panel) and iHP (right panel). **(C** and **D)** Individual examples of SWRs and replay that occurred during occupation of the reward zone. 150–250-Hz filtered LFP, color-coded reactivated spikes, and posterior probability of decoded position during an SWR are shown. The boundary of the SWR is indicated by a black dashed line; the calculated linear regression line is indicated by a red dashed line; and the statistical significance of permutation tests is denoted by an asterisk. **(E)** Pie chart showing the proportion of significant replay events between the dHP and iHP that occurred during occupation of the reward zone. The numbers of significant replays and candidate replay events are denoted in the bar. **(F)** The proportion of significant replays representing paths leading to either high-value or low-value reward zones. **p < 0.01.

However, in the iHP, no such robust spatial replays were observed. Instead, place cells in the iHP seemed to collectively reactivate spatial representations near either the intersection of the T-maze (ripples #1-4 in Figure 8D) or the choice arms (ripple #5-8 in Figure 8D). The proportion of significant replay events among candidate replay (see Methods for details) in the dHP (n = 28/185; 15.1%) was significantly higher than that in the iHP (n = 28/185; 15.1%) (p = 0.0019; two-sample Chi-square test) (Figure 8E). Importantly, most spatial replays in the dHP (n = 25/28; 89.3%) represented the rat’s navigational path leading to the high-value reward zone (p < 0.01; one-sample Chi-square test) (Figure 8F). This result aligns with previous spatial replay literature (Pfeiffer and Foster 2013, Michon et al. 2019). The robust spatial replay observed in the dHP also suggests that the lack of value-related changes in the reactivation patterns of place cells in the dHP compared to the iHP cannot be attributed to the idiosyncratic experimental conditions of the current study.

## Discussion

The current study investigated whether place cells in the iHP are reactivated differentially during SWR events after learning spatial value changes. We found that in the reward reversal block, high-value-coding place cells (HiV-cells) in the iHP were reactivated more frequently than low-value-coding place cells (LoV-cells) or location-coding place cells (Location-cells). Furthermore, selective reactivation of HiV-cells in the iHP was enhanced after rats learned the task after they acquired the reverse relationship between place and its associated reward value; moreover, this potentiated reactivation of HiV-cells in the iHP was maintained in the post-sleep session. Importantly, the frequent reactivation of HiV-cells in the iHP during the post-sleep session facilitated place-value learning on the next day. However, such value-based, learning-dependent reactivation was rarely observed in the dHP. Instead, we found that the spiking activities of place cells whose place fields were sequentially organized toward the high-value reward location were replayed more frequently in the dHP than in the iHP during SWRs. Overall, our findings suggest that functionally heterogeneous types of information are differentially and simultaneously reactivated in the dHP and iHP to optimize goal-directed navigation for maximal reward.

### Spiking activities representing high-value locations are selectively reactivated in the iHP but not in the dHP

In our experiments, we assumed that a subset of place cells that remapped their fields to represent high-value locations (HiV-cells) are specialized to encode the high value of a place. We found that HiV-cells in the iHP had higher reactivation probabilities than LoV-cells or Location-cells. These results suggest that the relative value of information associated with a place can modulate the reactivation probabilities of place cells in the iHP. Although some prior studies have suggested that the function of the iHP is to encode and update the value of a place, to the best of our knowledge, the type or quality of information reactivated in the iHP has rarely been previously reported (Bast et al. 2009, De Saint Blanquat et al. 2013, Jin and Lee 2021).

In our previous study (Jin and Lee 2021), we found that place cells in the iHP exhibited more robust remapping in response to changes in reward value than those in the dHP, and their fields accumulated near high-value locations. Another study by De Saint Blanquat et al. (2013) showed that inactivating the iHP by injection of muscimol impaired place-value updating in place-preference tasks, but no impairment was observed when the dHP was similarly inactivated. Furthermore, lesioning the iHP impaired the rapid learning of a place-learning task in which the location of the hidden escape platform changed daily (Bast et al. 2009) and abolished the goal-related activities of the medial prefrontal cortex in the place-preference task (Burton et al. 2009). These results imply that high-value–associated spatial information is reactivated selectively in the iHP and serves to facilitate associative learning between a place and its value.

In contrast, we found no such reactivation patterns in the dHP. Most studies of the dHP have focused on reinstatement of navigational trajectories (spatial replay) by sequential spatial firing (Widloski and Foster 2022, Mou et al. 2022, Igata et al. 2021, Gillespie et al. 2021, Bhattarai et al. 2020). However, few studies have directly investigated whether the reactivation patterns of cells are dependent on the relative values associated with different places in the dHP. It is worth noting that some study results imply that reward information can modulate the reactivation probabilities of place cells in the dHP. For example, Ambrose et al. (2016) showed that the frequency of reverse replay in the dHP was manipulated by the amount of reward provided in the reward zone in an experimental paradigm in which rats shuttled between two reward zones in a linear maze. Another study by Singer and Frank (2009) argued that reactivation responses were enhanced in the reward compared to the no-reward condition. These results are seemingly in conflict with other studies demonstrating that place cells in the dHP tend to exhibit value-insensitive firing when values associated with fixed reward locations changed (Jin and Lee 2021, Duvelle et al. 2019, Tabuchi et al. 2003). This discrepancy may reflect the difference in behavioral tasks; in this case, rats were required to update spatial value from a low-value (i.e., a small amount of reward) to a high-value (i.e., a large amount of reward) in a goal-directed navigational task (Jin and Lee 2021, Duvelle et al. 2019, Tabuchi et al. 2003). In contrast, other studies used non-hippocampal shuttling tasks (Ambrose et al. 2016) or provided rewards only in one of two reward zones in a W-shaped maze (Singer and Frank 2009).

Notably, some prior studies manipulated spatial value by associating an electric shock with a place (Moita et al. 2004). In this scenario, place fields in the dHP underwent global remapping after the animal experienced the electric shock. Our results are consistent with the idea that the spiking activities of place cells that remapped after associating a negative value with a certain place could also be reactivated more frequently than non-remapping cells. Ormond et al. (2023) tested this idea using an aversive spatial decision-making task in which rats received an electric shock in the choice arm. After experiencing the shock, rats did not re-enter the shock-associated arm, and a subset of place cells in the dHP remapped their fields globally. Importantly, reactivation probabilities were higher in place cells that exhibited shock-related remapping in the dHP than in non-remapping cells. In another study, rats received an electric shock at the end of a linear track (Wu et al. 2017). After the shock experience, rats avoided the shock zone as they moved around on the track, and the place cells whose fields were located in the shock zone in the pre-shock trials were reactivated more frequently during SWRs than those representing the non-shock zone in the pre-shock period. These results suggest that a negative event at a particular location changes the value of the place and may alter the probability of reactivation of the cells representing the location, serving to strengthen the associative memory between the place and its value during SWRs.

### Sequential spatial representations leading to the high-value location are replayed in the dHP but not in the iHP

Previous studies reported that spatial paths leading to goal locations are sequentially replayed more in the dHP than those leading to non-goal locations (Pfeiffer and Foster 2013, Michon et al. 2019, Igata et al. 2021). Based on our prior study that place cells in the iHP overrepresent high-value locations compared with low-value locations (Jin and Lee 2021), we originally expected spatial replays leading to goal locations to occur more frequently in the iHP than the dHP. However, sequential reactivation of spatial representations leading to high- and low-value locations was barely detectable in the iHP. Instead, we found robust spatial replays in the dHP, as has been frequently reported (Foster and Wilson 2006, Karlsson and Frank 2009, Diba and Buzsaki 2007). The iHP may not be an ideal region for the occurrence of such sequential replays of precise location information because the size of the place field is larger and spatial tuning is poorer in the iHP compared with the dHP (Jin and Lee 2021, Kjelstrup et al. 2008, Jung et al. 1994). Instead, the larger place fields in the iHP may make the area suitable for representing spatial values, given that the spatial resolution of value is expected to be lower than that of a place in a natural environment.

In the current study, 85% of replay events in the dHP represented sequential paths to the higher-value location, whereas only 15% of replay events represented paths to the low-value location. Thus, the direction of spatial replay in the dHP seems biased toward the high-value location, which aligns with prior studies well (Pfeiffer and Foster 2013, Michon et al. 2019, Igata et al. 2021). For example, using a dual-environment reward-place association task on an eight-arm maze, Michon et al. (2019) found that hippocampal replay events are biased toward paths associated with high-value (i.e., large reward) locations compared with those associated with low-value reward locations. Other studies reported that, when rats learned the location associated with reward in an open arena, trajectories leading to that location were replayed more frequently than those leading to no-reward locations (Pfeiffer and Foster 2013, Igata et al. 2021).

In addition, spatial replays could be directed to non-goal locations where animals underwent fearful events or where they received satiated type of reward. Specifically, Wu et al. (2017) identified an imaginary spatial replay leading to the place where rats underwent an electric shock at the end of linear track, even though rats did not enter the shock zone. Carey et al. (2019) used different motivational states (i.e., thirsty or hungry) to bias the rat’s navigation toward the choice arm containing the deprived reward (i.e., water or food) in the T-maze. They reported that replay occurred in the opposite direction relative to where the restricted reward was located, although rats shifted their behavior toward the restricted reward location. These results collectively suggest that the spatial replay of place cells in the dHP is modulated by spatial value information when different places are associated with positive or negative values in an environment.

### Reactivation of place cells in the iHP coding high spatial values is potentiated by subsequent sleep

It is well known that neural firing patterns acquired during a recent experience are reactivated in the hippocampus during subsequent sleep. For example, two place cells whose fields overlap on the open arena exhibit an increased tendency to be reactivated together during the following sleep session (Wilson and McNaughton 1994). Also, the firing sequence of place cells during active behavior is observed again during reactivation in the post-sleep session (Lee and Wilson 2002). In the current study, we found that place cells coding high spatial values in the place-preference task had higher reactivation probabilities during SWRs, and they sustained their potentiated reactivation during subsequent post-sleep sessions. The more frequent reactivation of motivationally salient information during sleep may facilitate the selective consolidation of such memories from the hippocampus to neocortical networks. Consistent with this idea, we found a positive correlation between the amount of sleep reactivation of HiV-cells in the iHP and learning speed the next day. These results imply that higher-value spatial events are replayed more during post-behavioral sleep, presumably for their high priority during consolidation, and that learning on the next day may depend on such selective consolidation.

## Acknowledgments

This study was supported by the National Research Foundation of Korea (NRF 2018R1A4A1025616, 2019R1A2C2088799, 2021R1A4A2001803, 2022M3E5E8017723, 2022R1I1A1A0106893511) and the BK21 FOUR supported by the Ministry of Education and NRF. We thank Joseph Shin for conducting some pilot experiments for this study and aiding in the collection of electrophysiological data.

## Author Contributions

J-S.W. designed the research, collected and analyzed the data, and wrote the manuscript. I.L. supervised all aspects of the research and wrote the manuscript.

## Materials and Methods

### KEY RESOURCE TABLE

Provided in a separate file

### LEAD CONTACT AND MATERIAL AVAILABILITY

Further information and requests for resources and materials should be directed to and will be fulfilled by the Lead Contact, Inah Lee.

#### Subjects

Six male Long-Evans rats weighing 300-400 g were used. Food was restricted to maintain body weight at 85% of free-feeding weight to sustain a higher motivation level in behavioral tasks; water was available ad libitum. Animals were housed individually under a 12-hour light/dark cycle. All protocols and procedures conformed to the guidelines of the Institutional Animal Care and Use Committee (IACUC) of Seoul National University.

#### Behavior paradigm

##### Place-preference task (T-maze)

An 8-arm maze (central platform size, 401 cm^2^; arm size, 8 × 45 cm; height, 70 cm from the floor) was converted to a T-maze by installing a T-shaped transparent acrylic structure in the center platform. The rat was confined to the start arm by an acrylic blocker. A food well (2 cm in diameter and 8 mm in depth) was located at the end of each arm, with a proximity sensor installed underneath the well to detect the displacement of an acrylic disc (2.5 cm in diameter) that covered the food well. An infrared sensor was installed at each arm entry. Transistor-transistor logic (TTL) sensor signals were transmitted to a data-acquisition system (Digital Lynx SX; Neuralynx). An LED-light complex to illuminate the experimental room and digital cameras to monitor animals’ positions were installed on the ceiling. The maze was surrounded by circular black curtains to which a considerable number of visual cues were attached. White noise (80 dB) was delivered through two loudspeakers during the recording session to mask unwanted noise in the environment. In the place-preference task, rats were required to choose an arm baited with a more preferred type of reward (i.e., sunflower seeds [SS] or whole sunflower seeds [4/4 SS]) over a less-preferred reward (i.e., Cheerios cereal [CR] or quarter sunflower seeds [1/4 SS]). Rats initially learned the arm in which the more preferred reward was baited by trial and error in Block 1. In Block 2, rats were started from the opposite start arm to make the task more hippocampal-dependent (Packard and McGaugh 1996). In Block 3, they started from the original starting arm but the locations associated with higher-value and lower-value rewards were reversed. This experimental design sought to examine the rat’s behavior and neural correlates when updating place and its associated value in the environment (Figure 1A). Each block was terminated when rats chose the arm baited with the preferred reward more than 12 times in the previous 15 trials. After the rats had been tested for 2 days with SS and CR rewards, they were tested in the same task, but this time were provided the preferred type of reward (SS) only in the two reward arms but in different amounts (i.e., 4/4 SS vs. 1/4 SS). The higher-value location on the first day was pseudo-randomly determined, and all rats departed from the south arm in Block 1. The location of the high-value location the next day was the same as that of the last high-value location the day before.

#### Maze pre-training and surgery

After rats had become familiarized with the maze environment for a few days (30 min/d), they were trained in a shuttling task in which they alternated between the two adjacent arms of the 8-arm maze to obtain rewards. Rats were trained to complete 240 trials in 1 hour, which took approximately 4 days (mean, 3.8 days; SD, 1.1 days). After pre-training, a hyperdrive containing 24 tetrodes was implanted for recording spikes from single units and local field potentials (LFPs). Tetrodes were made using platinum wires (17.8 μm in diameter). A semi-automatic plating device (Nano-Z; Neuralynx) was used to adjust the final impedance of each tetrode to approximately 130 kΩ (measured in gold solution at 1 kHz). The hyperdrive was configured to have two separate electrode bundles, one carrying 18 tetrodes to target the iHP and vHP (coordinates for implantation: 5 mm posterior to bregma, 6 mm lateral from midline), and the other containing 6 tetrodes to target the dHP (coordinates for implantation: 3.2 mm posterior to bregma, 3 mm lateral from midline). The bundles were inserted obliquely (5 degrees medially from the vertical axis) into the hyperdrive frame so that the tetrodes could maximally target the cell layers in the iHP.

#### Post-surgical training and main recordings

After 1 week of recovery from surgery, rats were retrained in the shuttling task. Retraining of rats to the same pre-surgical criterion required 10 ± 3 days (mean ± SD). During the post-training period, tetrodes were gradually lowered into the CA1 cell layer in the hippocampus while the rat was at rest on a pedestal. Once most tetrodes reached their target cell layers in the dorsal and intermediate-ventral regions in the hippocampus, the main recordings began. Place cell activity was recorded while rats shuttled on the two adjacent arms (spatial alternation task) for 7 days according to a previously detailed experimental protocol (Jin and Lee 2021). On days 8 to 11, rats performed the place-preference task in the T-maze. SS and CR were used as high-value and low-value rewards for recording on days 8 and 9, respectively, and 4/4 SS (high value) and 1/4 SS (low value) were utilized for days 10 and 11, respectively. Sleep sessions were recorded for approximately 30 minutes before and after the place-preference task while rats were at rest or asleep.

#### Electrophysiological recording procedures

After 1 week of recovery, each rat was placed on a custom-built pedestal outside the experimental room and habituated to rest for tetrode adjustment. Tetrodes were lowered individually to the cell layers over ∼2 weeks. Neural activity was amplified (1,000-10,000 times) and digitized (sampling frequency, 32 kHz; filtered at 600-6,000 Hz for spiking data and 0.1-1,000 Hz for LFP) using a Digital Lynx system (Neuralynx). The rat’s position and head direction were measured using an array of red and green LEDs attached to a custom headstage complex coupled to a preamplifier (HS-36; Neuralynx). A ceiling camera recorded LED positions and fed the signal to a frame grabber (sampling frequency, 30 Hz).

#### Histological verification of electrode tracks

After completion of the main recording sessions, tetrode-tip locations were marked by passing a weak electrical current through each tetrode (10 μA for 10 s). On the next day, the rat was sacrificed using an overdose of carbon dioxide (CO_2_) and then was perfused transcardially, first with phosphate-buffered saline (PBS) and then with a 4% (v/v) formaldehyde solution. Thereafter, the brain was removed and kept in a 4% v/v formaldehyde-30% sucrose solution at 4°C until it sank.

The brain was sectioned in the coronal plane at 40-μm thickness using a sliding microtome (HM 430; Thermo-Fisher Scientific), mounted on a slide glass, and then stained with thionin. Photomicrographs were taken using a digital camera attached to a microscope (Eclipse 80i; Nikon). Tetrode tracks were reconstructed using a series of sections, taking into account the presurgical configuration of the tetrode-array bundle and electrolytic lesion marks. Dorsoventral (DV) recording positions of place cells were quantitatively measured as the vertical distance from the cortical surface to electrode tip locations.

#### Data analysis

##### Unit isolation

Spikes associated with single units were isolated using a Windows-based, custom-written program (WinClust). The parameters, peak, valley, energy, and spike width, calculated from waveforms recorded from the four channels of a tetrode, were used for isolation. Unit-isolation quality was evaluated during cluster-cutting procedures, with each cluster defined as isolation quality 1 (poorly isolated) to 5 (well isolated) based on how well the cluster was separated from neighboring clusters and background noise. Units with an isolation rating of 1 were excluded from further analysis. Inter-spike interval (ISI) histograms were also used to judge the quality of the cut cluster. Units showing a mean firing rate > 10 Hz (either in the open field or in the radial maze) with a spike width < 300 μs were classified as inhibitory interneurons and were excluded from further analysis.

##### Sharp-wave ripple detection

SWRs were extracted from local field potentials (LFPs) using the following steps. First, the two tetrodes that recorded the largest SWRs were selected and the corresponding LFP signals were filtered with a 150-250 Hz bandpass filter. Sections exceeding 4 SDs from the average of the filtered signals were initially selected, and the point where enhanced signals decreased by less than 1 SD was taken as the boundary of the SWR for each tetrode. The SWR boundary was determined as the sum of the SWR boundaries of two tetrodes. If only one tetrode exceeded the threshold within a particular section, it was not counted as an SWR. Next, the number of cells reactivated within the SWR boundary was measured. If the number of reactivated cells was less than 5% of the total number of recorded cells, the signal was considered noise. Finally, signals recorded during passage of an SWR boundary were also considered noise if the animal’s head movement speed at the time was > 20 cm/s (e.g., bumping noise due to collision with the acrylic blocker installed in the T-maze). For spectrogram analysis, we selected the tetrode that exhibited maximal SWR amplitude in the dHP and iHP, respectively. We used the built-in Matlab function ‘mtspecgramc’ from Chronux 2.12 to calculate the spectrogram. The detailed parameters used were as follows: ‘fpass = [50 350]’, ‘tapers = [3 5]’, ‘trialave = 1’, ‘moving window = 100ms with 10ms increments within a window ranging from -200ms to +200ms’.

##### Basic firing properties

To construct a rate map, we first scaled the 720 × 480 pixel space down to 72 × 48 pixel space (1 pixel = 2 × 2 cm). The firing rate associated with a given pixel was calculated by dividing the number of visits by the number of spikes fired in each pixel. The raw rate map was then smoothed using an adaptive binning method. The amount of spatial information associated with a single spike of a single unit was measured by calculating spatial information based on the firing rate map, according to the following equation (Skaggs et al. 1992):

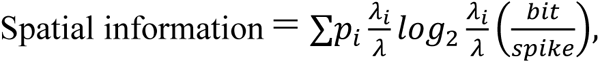

where *i* denotes bin, *p_i_* is the occupancy rate in the *i*^th^ bin, *λ_i_* indicates the mean firing rate in the *i*^th^ bin, and *λ* is the overall mean firing rate. Stability within the square box test was measured by comparing the firing rate map between the first half and the second half of the session using Pearson’s correlation.

##### Behavior analysis

For learning curve analyses, we calculated the estimated learning curve and the trial at which the rat reaches acquisition onset by applying a state-space model to behavioral performance (Smith et al. 2004). Latency is the time elapsed from passing by the infrared sensor installed at the entry of the start arm to reaching the food well. Learning speed is the inverse value of the trials to reach the learning criterion. In Figure 7F and 7H, we measured the learning speed in Block 1 on the next day. Because the high-value location of the first block on the day of the experiment was the same as the last block on the previous day, faster learning by rats in the first block was taken to mean that the information of high-value locations was strongly strengthened by sleep reactivation of high-value-coding place cells in the iHP.

##### Place cell analysis

A place cell was defined by applying the following steps. First, the spatial rate map, smoothed using Skaggs’s adaptive binning method, was calculated, and the pixel with the maximal firing rate was identified. Then, the firing rates associated with all other pixels were compared against the maximal firing rate, and only those pixels whose firing rate exceeded 20% of the peak firing rate were retained for further analysis. Among the remaining pixels, one or more sets of continuous pixels (>40 cm^2^ in size with a peak firing rate > 1 Hz) were defined as place fields. In cases where multiple place fields were found during the procedure, the field size of the unit was calculated by taking the sum of all subfields as the unit’s field size. Spiking data were included only if instantaneous speeds were greater than 5 cm/s at the time of the spike; spikes in the reward zones were excluded from the analysis (Lee et al. 2006). Because a proximity sensor was installed inside the food well, the exact timestamp when rats displaced the disc covering the reward could be measured. A cell was operationally defined as a place cell if 1) its field size was greater than 40 cm^2^, and its peak and mean firing rates were greater than 1 Hz and 0.25 Hz, respectively, and 3) its spatial information was higher than 0.25 bits/spike with a statistical significance of p < 0.01(Skaggs et al. 1992, Lee et al. 2004).

Place cells were divided into non-remapping and remapping place cells based on Pearson’s correlation coefficient (R) of spatial rate maps between Block 1 and Block 3. If R was greater than 0.75, it was regarded as a non-remapping place cell; otherwise, it was classified as a remapping place cell (Figure 3).

##### Reactivation analysis

Reactivation probability and reactivation rate were calculated using SWR events that occurred in Block 3, where location- and value-coding place cells were best dissociated. Reactivation probability was calculated by dividing the number of SWRs in which one place cell was reactivated by the total number of SWRs. Reactivation rate was measured by dividing the number of total reactivated spikes by the sum of total SWR durations. The same calculations were applied to pre- and post-sleep sessions, and the entire (∼30 min) pre- and post-sleep dataset was used for calculating reactivation activities.

##### Constructing the population rate map

Population rate maps for the place-preference task were constructed by linearizing spatial rate maps for all place cells and stacking them (bin size = 2 cm). Trials performed after reaching acquisition onset in Block 3 were used to create a spatial rate map, and the population rate map was created using Block 3. The firing rate in each bin was calculated by dividing the number of spikes by the number of occupancies by the rat within that bin. In the curved section, a fan-shaped boundary with an internal angle of 22.5 degrees was defined as a bin. Linearized firing rate maps were smoothed using a Gaussian window (window size, 22 cm; full width at half maximum [FWHM], 10 cm).

##### Bayesian decoding for position reconstruction

The rat’s positional information was reconstructed from the activity of place cells by applying a Bayesian decoding algorithm (Zhang et al. 1998, Brown et al. 1998). In Bayesian analysis, value-coding place cells, location-coding place cells, and place cells whose fields were located in the stem of the T-maze were pooled. Briefly, the number of spikes at a given time ***t***, *n_i_*(***t***), was calculated for each place cell, *i*. The 2D spatial rate map was linearized to a 1D spatial rate map and the mean firing rate at given spatial position ***x***, **λ***_i_*(***x***), was calculated. The reconstruction was obtained using two vectors, ***n***(***t***) = (*n*_1_, *n*_2_, … *n*_N_) and **λ**(***x***) = (*λ*_1_, *λ*_2_, … *λ*_N_), where ‘N’ is the number of recorded place cells and the firing probability was assumed to follow a Poisson distribution.

The animal’s position was reconstructed using the standard formula for conditional probability (1):

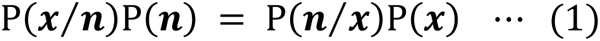

where P(*x*) is the probability that the animal is positioned at ***x***, and P(*x*/*n*) is the conditional probability that the animal is located at position ***x*** when the spike count was ***n***. This equation (1) is rearranged to (2) to solve for conditional probability of reconstructed animal’s position.

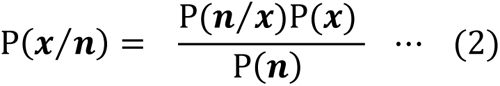

Assuming that the firing characteristics of each place cells follow a Poisson distribution and that all cells fire independently, the prior template, P(*x*/*n*), was calculated as (3):

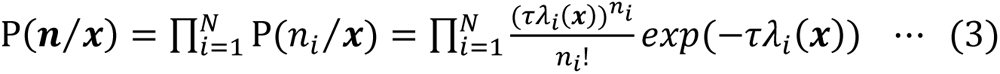

where τ is the time window for data sampling (20 ms). Thus, P(*x*/*n*) was calculated according to the equation (4):

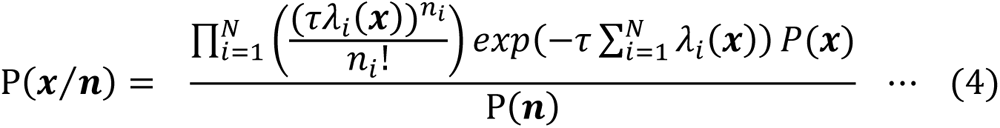

and P(*n*) was used to normalize probabilities (Zhang et al., 1998). The position at which P(*x*/*n*) was maximum was considered to be the reconstructed position (*x̂*):

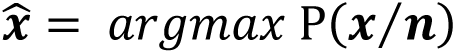

### QUANTIFICATION AND STATISTICAL ANALYSIS

Statistical testing was performed with MATLAB using built-in and custom-made functions. The null hypothesis was rejected using the non-parametric Wilcoxon rank-sum and Kruskal-Wallis tests. A Chi-square test was utilized to test the proportional difference between groups. The significance of linear relationships between two dependent variables was tested using linear regression. A Bonferroni correction was used to correct the significance level. Statistical results are reported in the Results sections. All tests were two-tailed, and significance was accepted at a p-value of 0.05.

## DATA AND CODE AVAILABILITY

Data and MATLAB code are available from the authors upon reasonable request.

---

**Figure S1.**
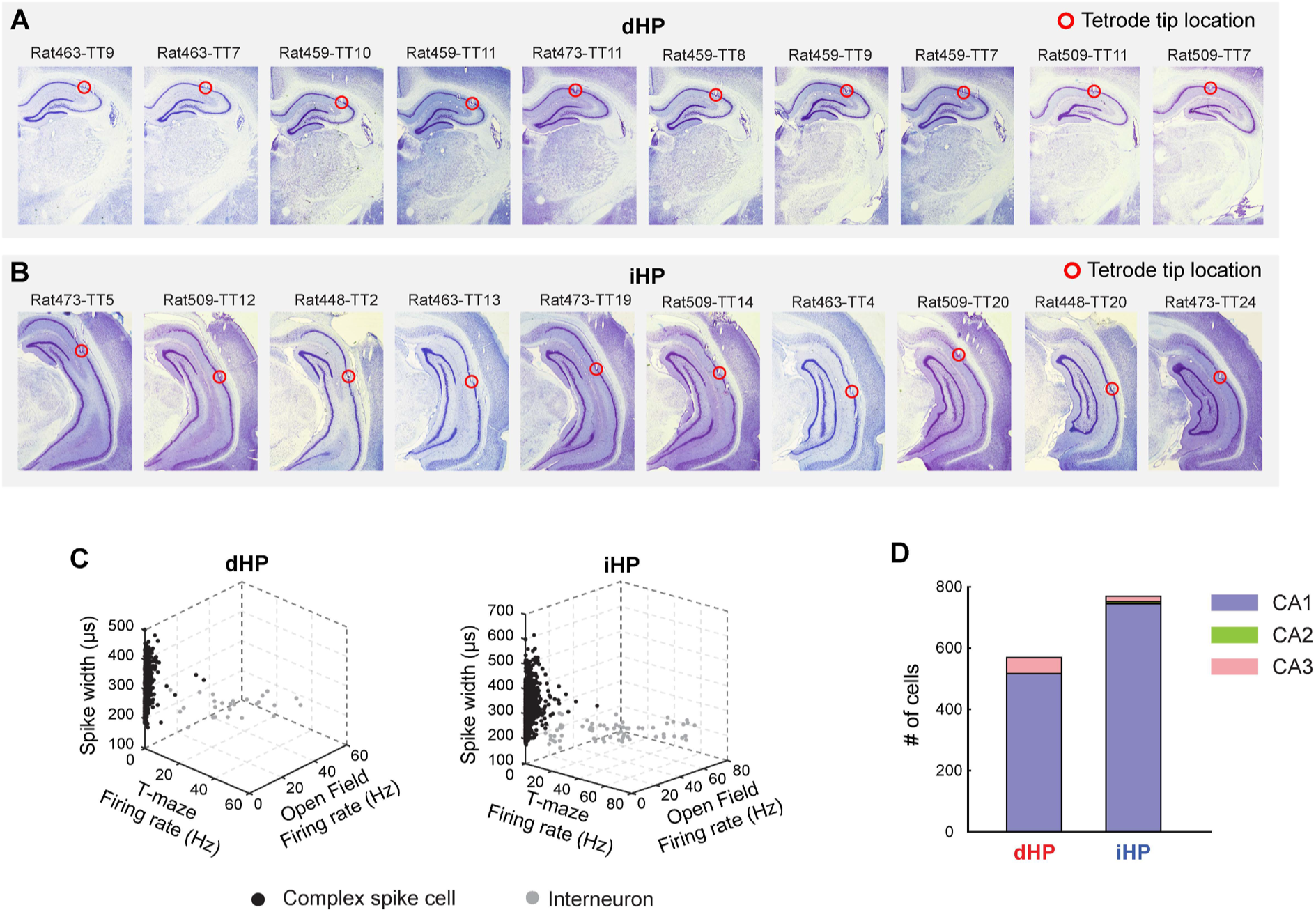
Histological verification of electrophysiological recording and basic information of recorded cells, related to Figure 2. **(A** and **B)** Histological verification of electrode tip locations where electrophysiological data were recorded across the dorsal (A) and intermediate (B) hippocampus. **(C)** Complex spike cells were distinguished from inhibitory interneurons by using the spike width and mean firing rates in the T-maze and open fields. **(D)** Stacked bar graphs showing which region of the hippocampal subregion the place cell was recorded in the dHP, iHP, and vHP.

**Figure S2.**
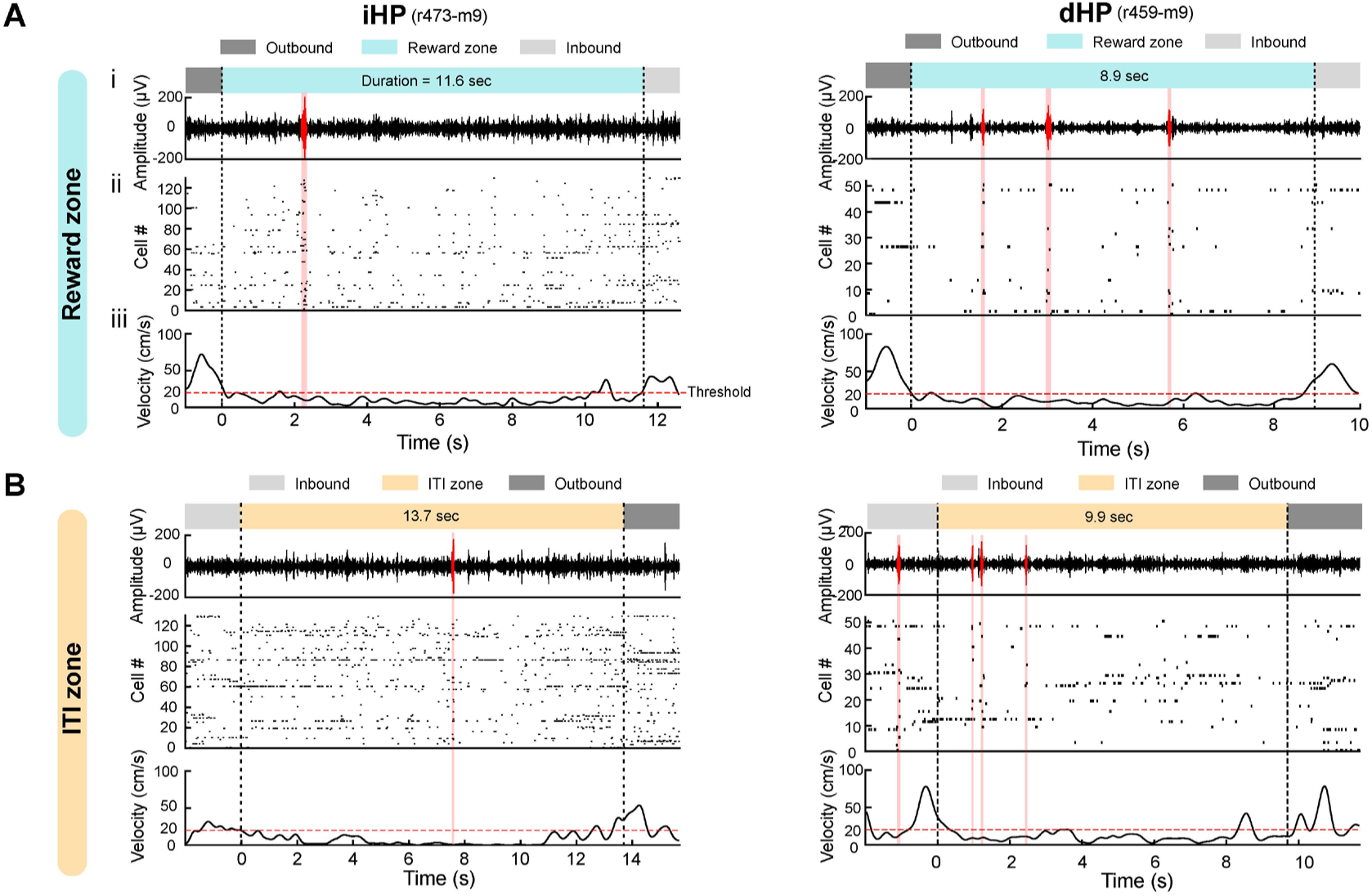
Examples of SWR detection during reward zone and inter-trial interval in iHP and dHP, related to Figure 2. **(A** and **B)** SWRs were detected when animals were in the reward zone (A) and ITI (B). The black dashed line indicated the boundary between behavioral periods. SWR should satisfy the following criteria of the large amplitude of 150 ∼ 250 Hz filtered LFP (A-i), enough ensemble activity (A-ii), and low movement speed (A-iii). SWR periods were indicated by the red shaded area. The examples of iHP and dHP were from rat 473 and rat 459, respectively. While rats were located in the reward zones, their velocity measured by customized LED headstage ranged from 0 to 20cm/s due to their head movement. By examining raw LFP, we confirmed that SWRs occurred when rats were in not only immobile status (< 5 cm/s) but also when their head was slightly moved without changes of body position (5∼20 cm/s). Thus, periods with speeds lower than 20 cm/s were considered immobile periods.

**Figure S3.**
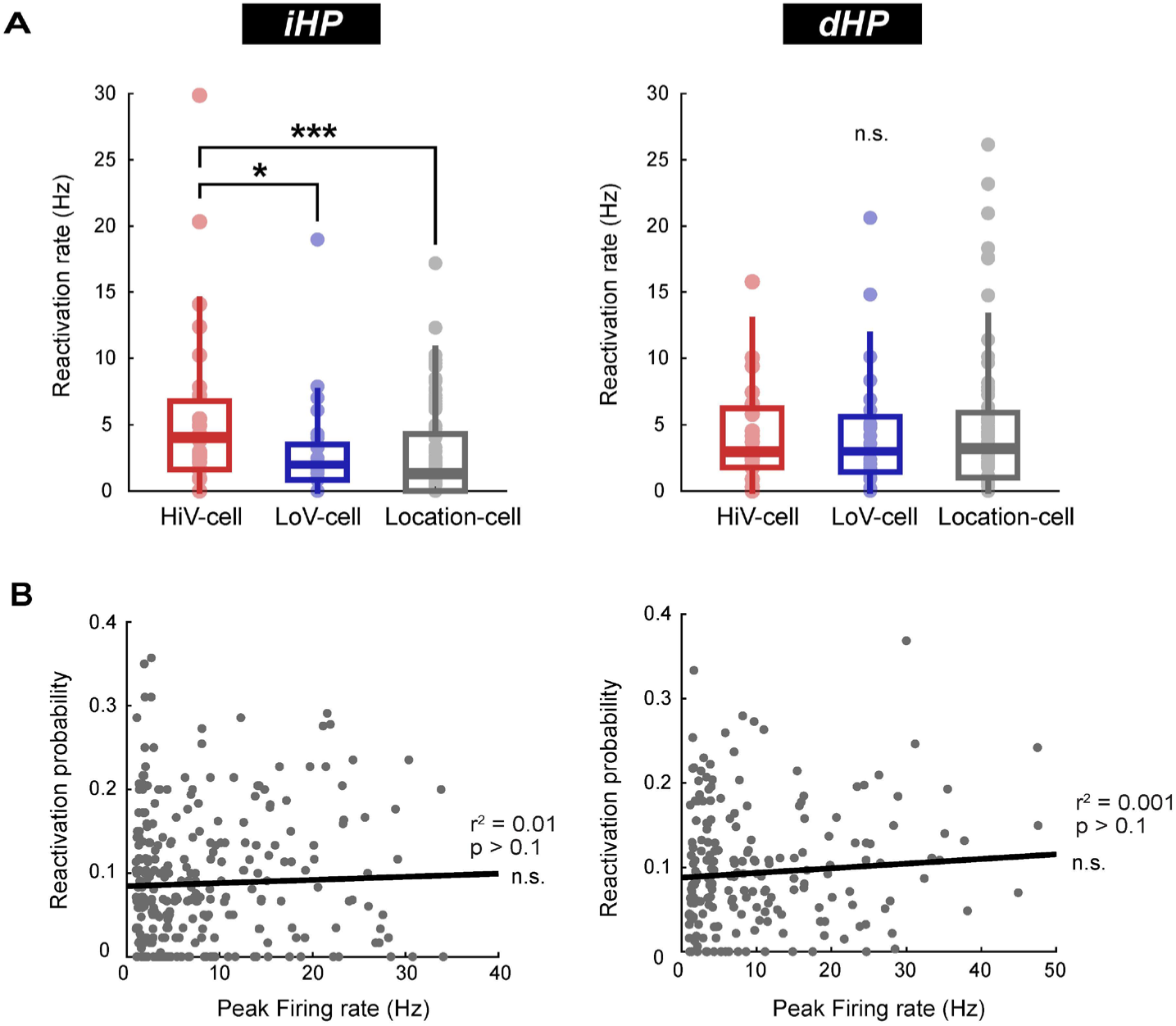
Alternative measurement showing exclusive reactivation of RMHi during SWR in the iHP but not in the dHP, related to Figure 4. **(A)** Comparison of reactivation rate in the iHP (left panel) and dHP (right panel). It is calculated by dividing the number of total reactivated spikes by the sum of SWR duration. Each dot indicated the reactivation probability of a single cell. **(B)** Cross-correlation between reactivation probability and peak firing rate of spatial rate maps to examine whether cells with higher firing rate reactivated more significantly than those with lower firing rate in the iHP (left panel) and dHP (right panel). Asterisks indicate statistical significance (*p < 0.05, ***p < 0.01).

**Figure S4.**
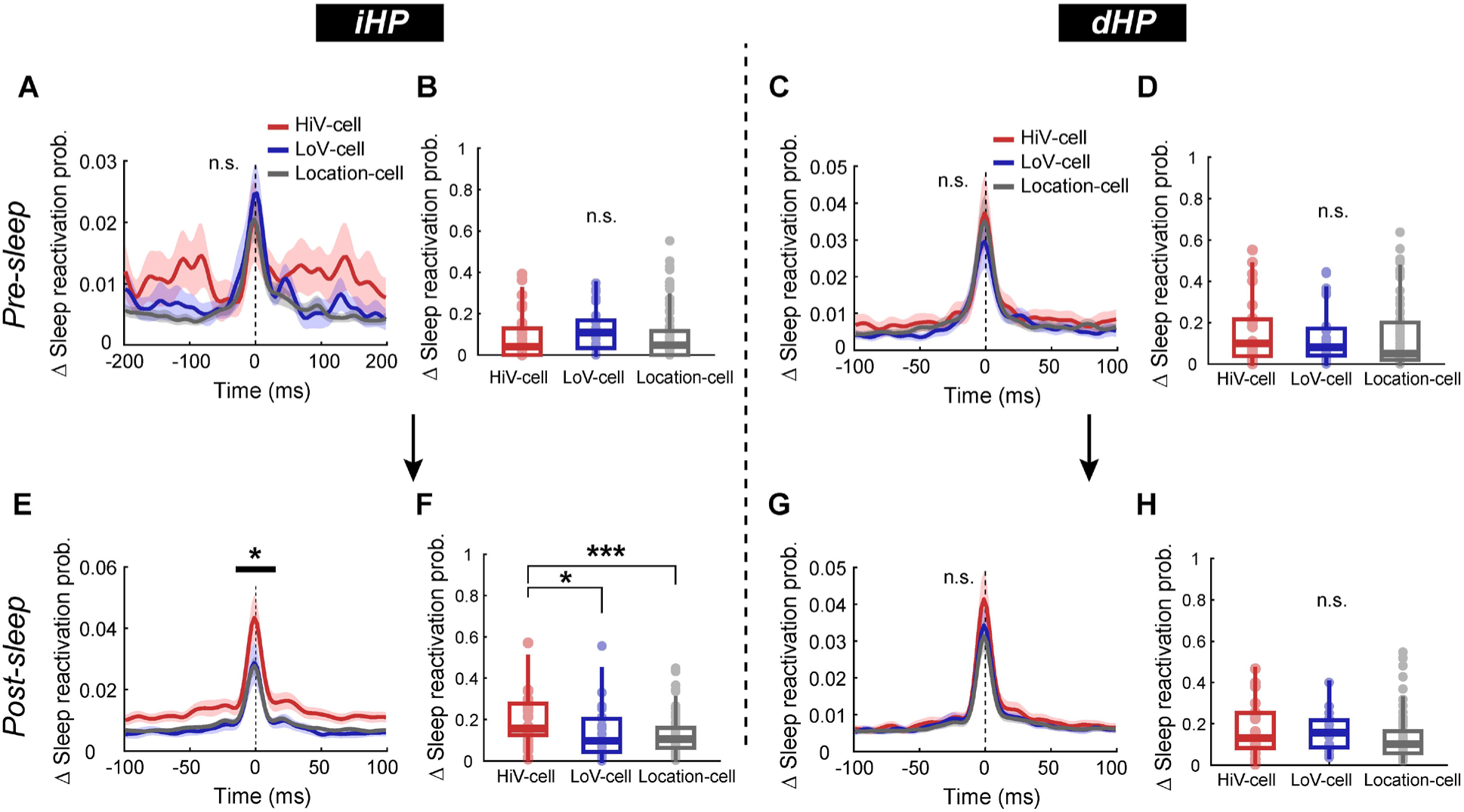
Separate analysis of reactivation patterns in pre- and post-sleep, respectively, related to Figure 7. **(A)** Peri-event time histogram aligning with the SWR peak time, illustrating the relationship between pre-sleep reactivation probability and SWR events depending on the cell types. **(B)** Comparison of pre-sleep reactivation probabilities in the iHP. Each dot denotes the reactivation probability of a single cell. **(C** and **D)** The same analyses as in (A and B) were replicated in the dHP. **(E – H)** The same analyses as in (A – D) were replicated in the post-sleep session. Asterisks indicate statistical significance (#p < 0.1, *p < 0.05, ***p < 0.001).

**Figure S5.**
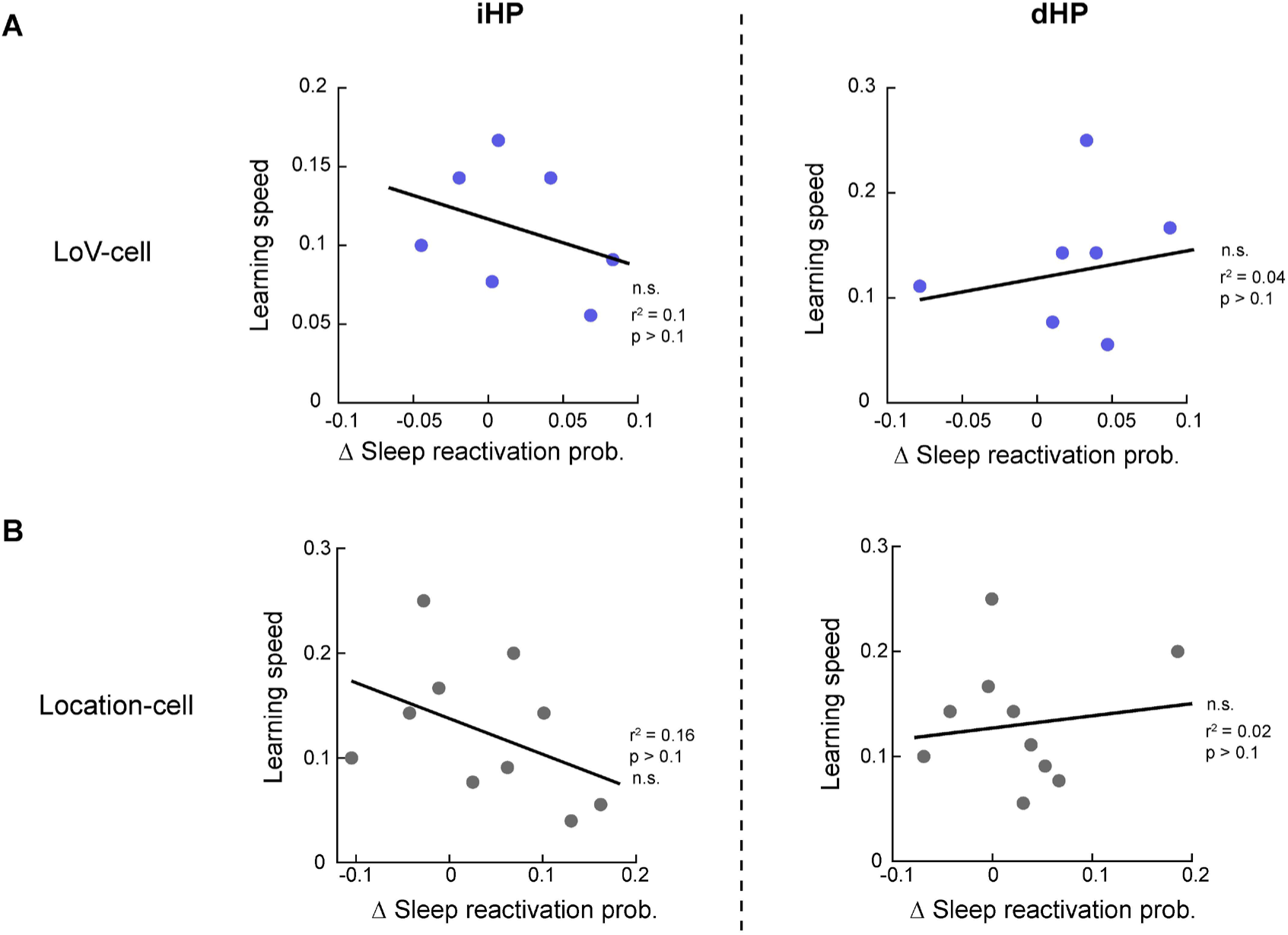
Relationships between learning speed on following day and inter-sleep reactivation difference of Low-cell and Location-cell, related to Figure 7. **(A** and **B)** Cross-correlation between learning speed on subsequent day and inter-sleep reactivation difference of Low-cell (A), Location-cell (B) were calculated based on session-mean reactivation probability in the iHP (left panel) and dHP (right panel). Each dot represented an average of reactivation probabilities per session.

